# Expanding the repertoire of chemically induced covalent neoantigens

**DOI:** 10.1101/2025.07.08.663715

**Authors:** Chen Zhou, Wei Huang, Xiaokang Jin, Haofeng Wu, Lynn M. McGregor, Markus Schirle, Chenlu Zhang, Xiaoyu Zhang

## Abstract

The ubiquitin-proteasome system generates peptide fragments that are displayed on MHC-I molecules. Recent studies revealed that covalent small molecules can hijack this process, leading to the presentation of hapten-modified peptide fragments, or covalent neoantigens. Here, we report a chemical proteomics platform to systematically investigate this process and harness covalent neoantigens for immune cell recruitment and activation. Using a bioorthogonal cysteine-reactive probe (IA-mTz), we demonstrate that diverse intracellular proteins can be modified, processed, and presented as covalent neoantigens across multiple cell lines. Mass spectrometry confirmed the presence of IA-mTz-modified peptides within the MHC-I immunopeptidome, and structural modeling revealed stable peptide-MHC-I interactions. We further integrated a bioorthogonal strategy that enables immune engagement: IA-mTz-modified cells conjugated with trans-cyclooctene (TCO)-IgG elicited Fc receptor activation in a reporter assay, supporting immune recruitment via antibody-dependent cellular phagocytosis. This approach was extended to additional cysteine-reactive probes and to 2,4-dinitrochlorobenzene (DNCB), a common skin sensitizer, which we show induces covalent neoantigen formation in a cell- and *HLA*-dependent manner. These findings reveal a previously underappreciated breadth of covalent neoantigen formation and provide a generalizable strategy for immune targeting of covalent neoantigens.

## Introduction

The ubiquitin-proteasome system (UPS) plays a crucial role in breaking down intracellular proteins into peptide fragments, some of which are presented by class I major histocompatibility complex (MHC-I) as peptide antigens^1, 2^. Recent studies have demonstrated that targeting intracellular proteins using covalent small molecules can interfere with the antigen processing pathway, resulting in the presentation of hapten-modified peptide fragments, or covalent neoantigens, on MHC-I^3, 4^. These unnatural epitopes create new opportunities for developing antibodies that specifically recognize them. Such antibodies can be integrated into bispecific T-cell engagers (BiTEs), which recruit cytotoxic T lymphocytes (CTLs) for the targeted elimination of cancer cells. One of the key advantages of this approach is its ability to target proteins regardless of their functions and mutational status. By modifying proteins, including those that are non-essential and non-mutant, cells have the potential to present covalent neoantigens that can be therapeutically exploited.

The three reported targets (KRAS-G12C, EGFR-T790M, and BTK) are all modified by their respective covalent ligands through engagement with cysteine residues^3, 4^. Across the human genome, more than 60,000 cysteines (a number that continues to grow) have been identified as reactive and susceptible to modification by electrophilic small molecules and metabolites^5, 6^. Given this vast reservoir of chemically modifiable cysteines, we asked whether covalent neoantigens might arise more broadly across the proteome and whether they could be harnessed for therapeutic applications or play roles in physiological and disease processes. In this work, we developed chemical probes and an integrated platform to investigate this concept. We found that covalent neoantigens can arise from a diverse array of peptide antigens across multiple cell lines and can be leveraged to develop new therapeutic modalities that recruit and activate immune cells. Finally, we demonstrated that 2,4-dinitrochlorobenzene (DNCB), a potent skin sensitizer, induces the formation of covalent neoantigens, providing a framework to explore the mechanisms underlying chemically induced allergic and immune responses.

## Results

### Development of a cell-permeable probe to identify covalent neoantigens

To investigate whether covalent neoantigens arise from a broad range of MHC-I-associated peptide antigens, we first aimed to develop a cell-permeable, broad-spectrum cysteine-reactive probe capable of modifying a wide range of intracellular proteins and assessing whether the modified proteins are ultimately processed and presented as covalent neoantigens on the cell surface (**Figure 1A**). We selected iodoacetamide as the cysteine-reactive electrophilic group due to its widespread use in profiling reactive cysteines, particularly within well-established activity-based protein profiling (ABPP) platforms^7, 8^. To enable detection of covalent neoantigens, we selected a bioorthogonal methyltetrazine (mTz) moiety to generate iodoacetamide-mTz (IA-mTz) (**Figure 1B**), which can be rapidly conjugated to trans-cyclooctene (TCO)-bearing compounds^9^ for downstream detection of IA-mTz-modified peptide antigens. IA-mTz exhibited cytotoxicity in the test cell line BV173, with an IC_50_ of 0.22 μM (**Figure S1A**), indicating efficient cell permeability and engagement with intracellular proteins, potentially inducing stress and apoptosis^10^. To further confirm its cell permeability, we treated both live cells and cell lysates with IA-mTz, followed by lysis and bioorthogonal ligation with biotin-TCO, then analyzed the extent of proteome labeling via Western blot for biotin. The results confirmed that IA-mTz effectively labeled proteins in live cells (**Figure S1B**).

**Figure 1.**
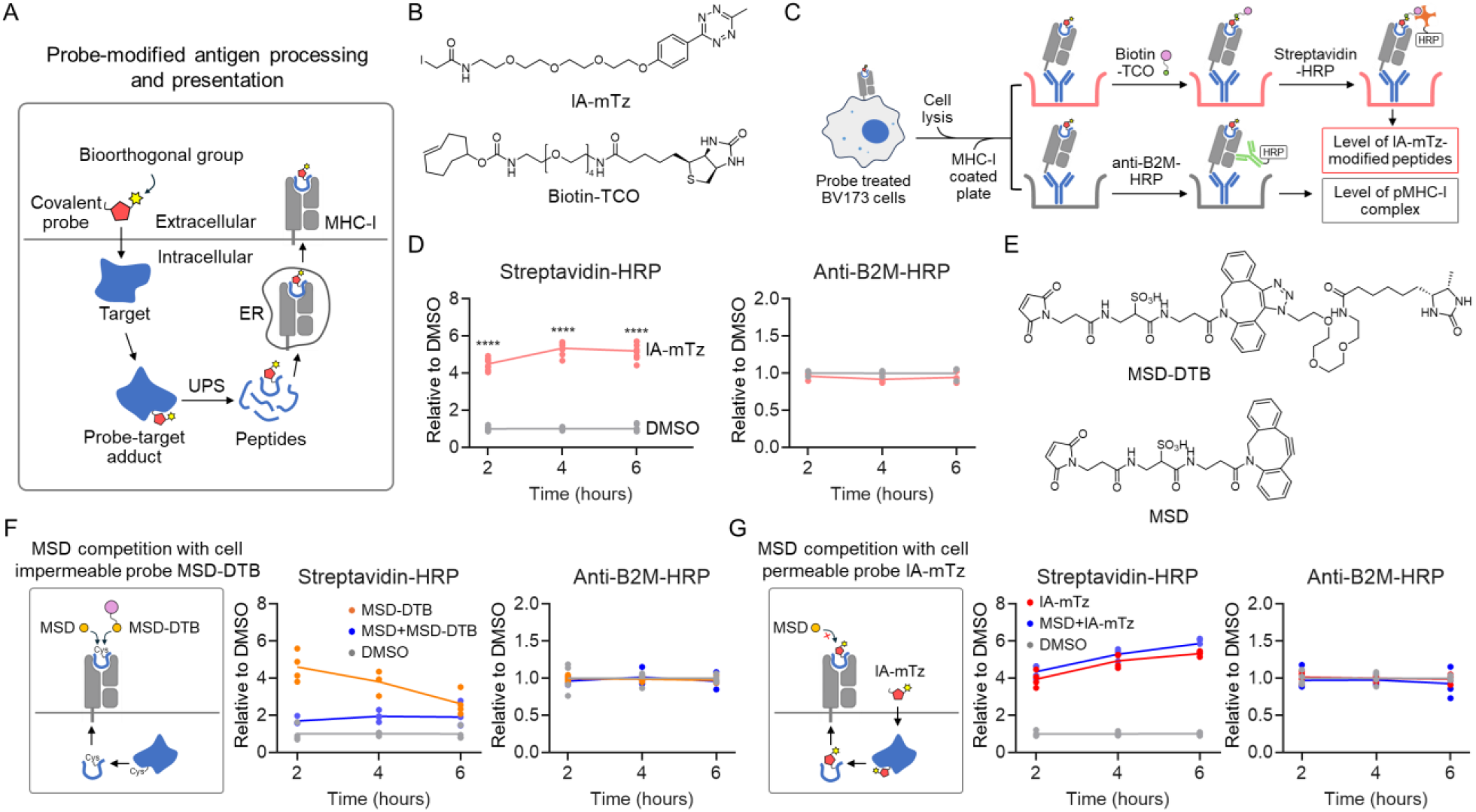
Development of a cell-permeable probe to identify covalent neoantigens presented on MHC-I. **A**. A schematic illustrating the use of covalent probes bearing a bioorthogonal handle to label intracellular proteins, interfere with antigen processing, and induce the formation of covalent neoantigens. **B**. Structures of iodoacetamide methyltetrazine (IA-mTz) and biotin-trans-cyclooctene (biotin-TCO). **C**. Schematic representation of the ELISA-based workflow for detecting and quantifying IA-mTz-modified peptide antigens presented on MHC-I. **D**. ELISA assay measuring IA-mTz-modified peptides and B2M levels within the pMHC-I complex. Data are presented as mean ± SEM (n = 4 independent replicates for streptavidin-HRP; n = 3 for anti-B2M-HRP). The statistical significance was assessed using unpaired two-tailed Student’s t-tests. **E**. Structures of MSD-DTB and MSD. **F**. ELISA assay measuring MSD-DTB-modified peptides and B2M levels within the pMHC-I complex. Data are presented as mean ± SEM (n = 4 independent replicates for streptavidin-HRP; n = 3 for anti-B2M-HRP). The statistical significance was assessed using unpaired two-tailed Student’s t-tests. **G**. ELISA assay measuring IA-mTz-modified peptides and B2M levels within the pMHC-I complex. Data are presented as mean ± SEM (n = 4 independent replicates for streptavidin-HRP; n = 3 for anti-B2M-HRP). The statistical significance was assessed using unpaired two-tailed Student’s t-tests.

### IA-mTz-modified intracellular proteins are processed and presented as covalent neoantigens

Next, we sought to determine whether IA-mTz treatment leads to the presentation of IA-mTz-modified peptide antigens on MHC-I. BV173 cells were treated with 0.25 µM IA-mTz for various time points (2, 4, and 6 hours), then washed, harvested, lysed, and incubated with an enzyme-linked immunosorbent assay (ELISA) plate coated with the pan-MHC-I antibody (clone W6/32) to enrich peptide-MHC-I (pMHC-I) complexes. The enriched samples were then subjected to two measurements: one portion was conjugated with biotin-TCO, followed by detection with streptavidin-horseradish peroxidase (HRP) to quantify IA-mTz-modified peptides within the pMHC-I complexes; the other was incubated with anti-β2-microglobulin (B2M)-HRP to assess total pMHC-I levels (**Figure 1C**). Results showed that IA-mTz treatment led to a significant increase in IA-mTz-modified antigens at all three time points without affecting total pMHC-I levels (**Figure 1D**).

A key question raised by this observation was whether the detected signal originated from intracellular protein modification followed by antigen processing and presentation, or from direct labeling of cell surface proteins. In our previous study^11^, three iodoacetamide-based probes showed negligible cell surface labeling in BV173 cells even at high concentrations (50 µM) with short treatment times (30 minutes), suggesting the likelihood that the observed signal in this study originates intracellularly. To further confirm this mechanism, we performed a blocking experiment. Cells were pretreated with a high concentration (50 µM) of the cell-impermeable probe maleimide-sulfonate-dibenzocyclooctyne (MSD) (**Figure 1E**) to occupy cysteines within the MHC-I immunopeptidome already present on the cell surface. Subsequently, cells were treated with either MSD-DTB or IA-mTz (**Figure 1F,G**). We hypothesized that MSD pretreatment would block labeling by the cell-impermeable MSD-DTB but not affect cell-permeable IA-mTz labeling, as the latter targets newly processed antigens derived from intracellular proteins. Indeed, MSD pretreatment drastically reduced MSD-DTB labeling of MHC-I antigens but had no effect on IA-mTz-mediated labeling (**Figure 1F,G**), confirming that IA-mTz labels intracellular proteins that are subsequently processed and presented as covalent neoantigens on the cell surface. We further tested IA-mTz in seven additional cell lines (A549, H1975, HCT116, HeLa, AsPC-1, 22Rv1, and MDA-MB-231) and observed a significant increase in IA-mTz-modified pMHC-I complexes in all cases (**Figure S2**), suggesting that this mechanism is broadly applicable across diverse cellular contexts.

### Identification of IA-mTz-modified peptide antigens within the MHC-I immunopeptidome

Next, we employed immunopeptidomics to identify IA-mTz-modified cysteine-containing peptide antigens within the MHC-I immunopeptidome. BV173 cells were treated with IA-mTz and thoroughly washed with phosphate-buffered saline (PBS) prior to harvesting to remove any residual free probe that could cause artifact labeling during lysis. Cells were then lysed, and pMHC-I complexes were enriched via immunoprecipitation using the pan-MHC-I antibody. The eluted immunopeptidome was analyzed by mass spectrometry (MS) (**Figure 2A**). 2,204 unique 8-12-mer peptides were identified, with a dominant 9-mer distribution pattern (**Figure 2B** and **Table S1**), consistent with previous reports^12^. Motif analysis of the 9-mer peptides showed strong alignment with the binding preferences of the BV173 *HLA* alleles (*HLA-A*\*02:01, *HLA-A*\*30:01, *HLA-B*\*15:10, *HLA-B*\*18:01, *HLA-C*\*03:04, and *HLA-C*\*12:03)^13^, suggesting successful enrichment of the MHC-I immunopeptidome (**Figure 2C**).

**Figure 2.**
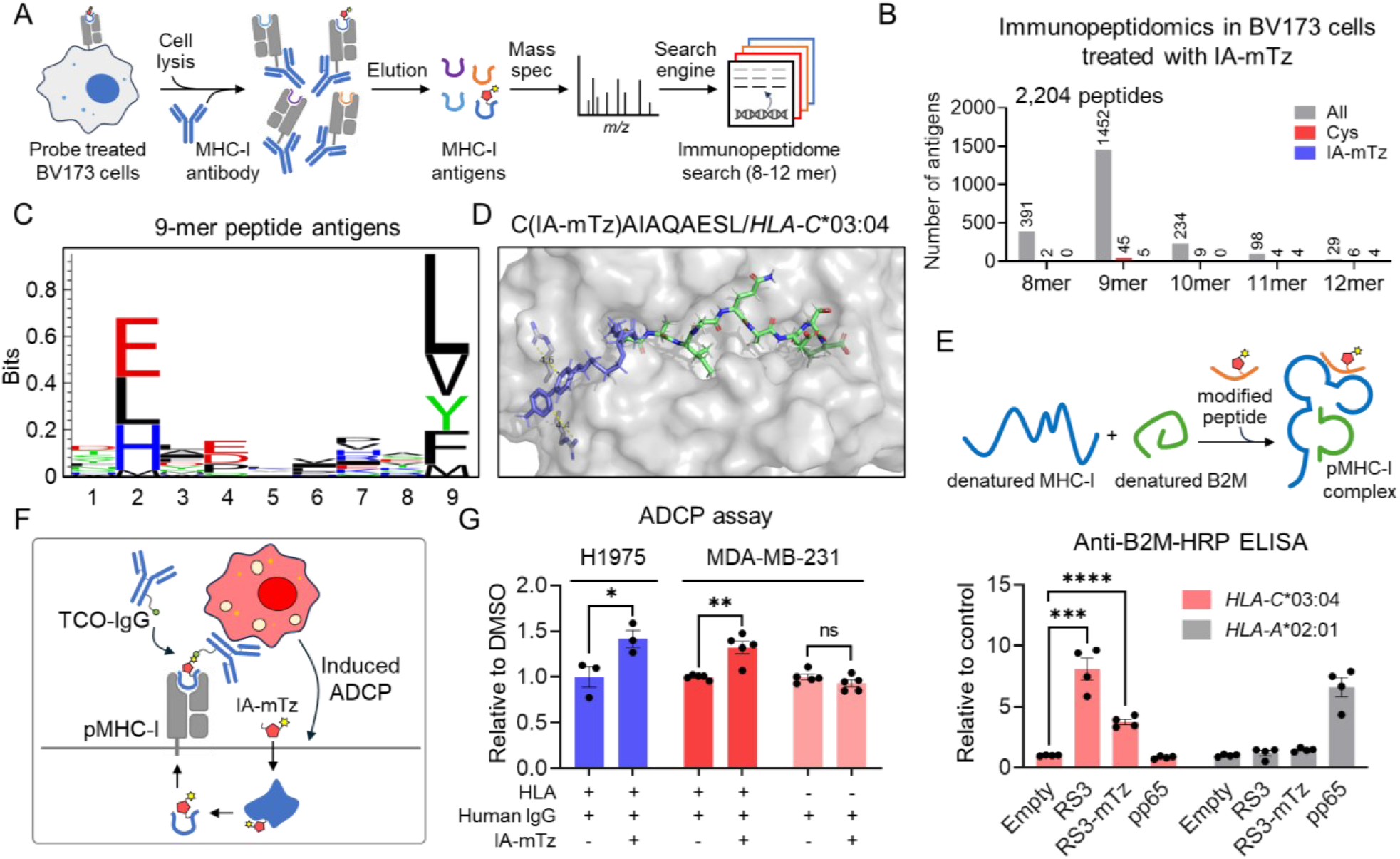
Identification of IA-mTz–modified peptide antigens within the MHC-I immunopeptidome and harnessing covalent neoantigens for immune cell recruitment and activation. **A**. Schematic representation of the immunopeptidomics experimental workflow. **B**. Distribution of 8-12-mer MHC-I-bound peptides. The result is representative of two independent experiments (n = 2). **C**. Motif analysis of all 9-mer MHC-I-bound peptide antigens. **D**. A modeling study suggests that the IA-mTz-modified RS3_97-105_ peptide fits within the binding groove of *HLA-C*\*03:04. **E**. *In vitro* MHC-I refolding assay assessing the association of unmodified and IA-mTz-modified RS3_97-105_ peptides with *HLA-C*\*03:04 and *HLA-A*\*02:01. The pp65 peptide (NLVPMVATV) serves as a control that binds *HLA-A*\*02:01 but not *HLA-C*\*03:04. Data are presented as mean ± SEM (n = 4 independent replicates). The statistical significance was assessed using unpaired two-tailed Student’s t-tests. **F**. Schematic representation of leveraging bioorthogonal chemistry and covalent neoantigens to induce ADCP. **G**. ADCP assay measuring luciferase activation in Jurkat-Luc NFAT-CD32 cells following co-culture with TCO-IgG and IA-mTz-pretreated target cells. Data are presented as mean ± SEM (n = 3 independent replicates for H1975; n = 4 independent replicates for MDA-MB-231). The statistical significance was assessed using unpaired two-tailed Student’s t-tests.

Among these, we identified five IA-mTz-modified 9-mer peptides (**Figure S3** and **Table S1**). We suspect the low number of modified peptides is due to the bulky size of the IA-mTz probe, which may reduce ionization efficiency, as well as the presence of a PEG linker that could fragment unpredictably, resulting in ambiguous MS1 and/or MS2 spectra. Nevertheless, we used the T cell epitope prediction tool from the Immune Epitope Database (IEDB)^14^ to assess MHC-I binding affinity of the five IA-mTz-modified peptides. One peptide antigen, RS3_97-105_ (CAIAQAESL), showed strong predicted binding to *HLA-C*\*03:04 in its unmodified form (**Figure S3**). Modeling of the IA-mTz-modified RS3_97-105_ peptide within the *HLA-C*\*03:04 binding groove revealed that the modification introduces additional hydrogen bonds with two residues in the MHC-I protein (**Figure 2D**). To biochemically validate these findings, we synthesized both unmodified and IA-mTz-modified RS3_97-105_ peptides and performed an *in vitro* MHC-I refolding assay^15^ to evaluate their ability to associate with MHC-I and B2M, forming a stable ternary complex measurable by ELISA (**Figure 2E**). The results revealed that both forms of the peptide could form stable complexes with *HLA-C*\*03:04, but not with *HLA-A*\*02:01 (**Figure 2E**). These findings provide direct evidence for the presence of IA-mTz-modified peptide antigens within the MHC-I immunopeptidome.

### Harnessing covalent neoantigens to recruit immune cells

The feasibility of leveraging covalent neoantigens to develop antibodies that recruit and activate T cells has been demonstrated in the context of three oncoproteins^3, 4^. Building on this concept, we sought to leverage our platform to expand this strategy through bioorthogonal chemistry. Specifically, we designed a strategy in which a trans-cyclooctene (TCO) moiety is conjugated to a second component capable of recruiting immune effector cells. The rapid bioorthogonal ligadation between mTz and TCO allows us to assess whether the resulting proximity between cells could further trigger immune activation. To this end, we conjugated TCO to human immunoglobulin G (IgG) (**Figure S4**). The resulting TCO-IgG can rapidly react with IA-mTz-modified peptide antigens displayed on the cell surface, therefore creating an IgG-coated cell state. The Fc region of IgG can then engage Fc receptors such as CD32 on macrophages or CD16 on natural killer (NK) cells^16^, leading to immune cell activation and clearance of the antibody-coated targets^17^.

To evaluate this strategy, we employed a Jurkat-Luc NFAT-CD32 reporter cell model, in which antibody-dependent cellular phagocytosis (ADCP) and subsequent luciferase expression are induced upon CD32 activation through interaction with IgG-coated cells (**Figure 2F**). Cells were treated with IA-mTz, washed to remove unbound probe, and incubated to allow antigen processing and presentation of IA-mTz-modified peptides via MHC-I. These cells were then co-cultured with Jurkat-Luc NFAT-CD32 cells in the presence of TCO-IgG, and the ADCP response was measured. In both H1975 and MDA-MB-231 cell lines, IA-mTz treatment led to a significant increase in ADCP activity (**Figure 2G**). To confirm that this response was mediated by pMHC complexes, we tested IA-mTz in MDA-MB-231 *HLA* knockout cells^11^ and observed negligible ADCP activation (**Figure 2G**). These results support that the ADCP response is driven by IA-mTz-modified peptide antigens presented via MHC-I. Collectively, these findings highlight the therapeutic potential of covalent neoantigens for immune recruitment and activation. Moreover, we present a novel strategy that integrates covalent neoantigen presentation with bioorthogonal chemistry to facilitate immune cell engagement.

### Expanding covalent neoantigens with additional chemical probes

We sought to determine whether other covalent probes could similarly induce covalent neoantigens and be leveraged to recruit and activate immune cells via the strategy described above. To this end, we selected two broad-spectrum scout fragments, KB02 and KB05, previously shown to bind a large portion of the human proteome and explore protein ligandability^7, 18^, and synthesized two analogs incorporating an mTz group (KB02-mTz and KB05CAA-mTz, **Figure 3A**). We confirmed that both probes are cell-permeable, as evidenced by intracellular proteome labeling (**Figure S5A**), and exhibited cytotoxicity at low micromolar concentrations (**Figure S5B**). Using ELISA assays, we observed the presentation of probe-modified peptide antigens within the MHC-I immunopeptidome (**Figure 3B**). Furthermore, ADCP assays demonstrated that these probe-modified bioorthogonal antigens on the cell surface effectively conjugate with TCO-IgG under physiological conditions, inducing ADCP activity in MDA-MB-231 parental cells, but not in *HLA* knockout cells (**Figure 3C**).

**Figure 3.**
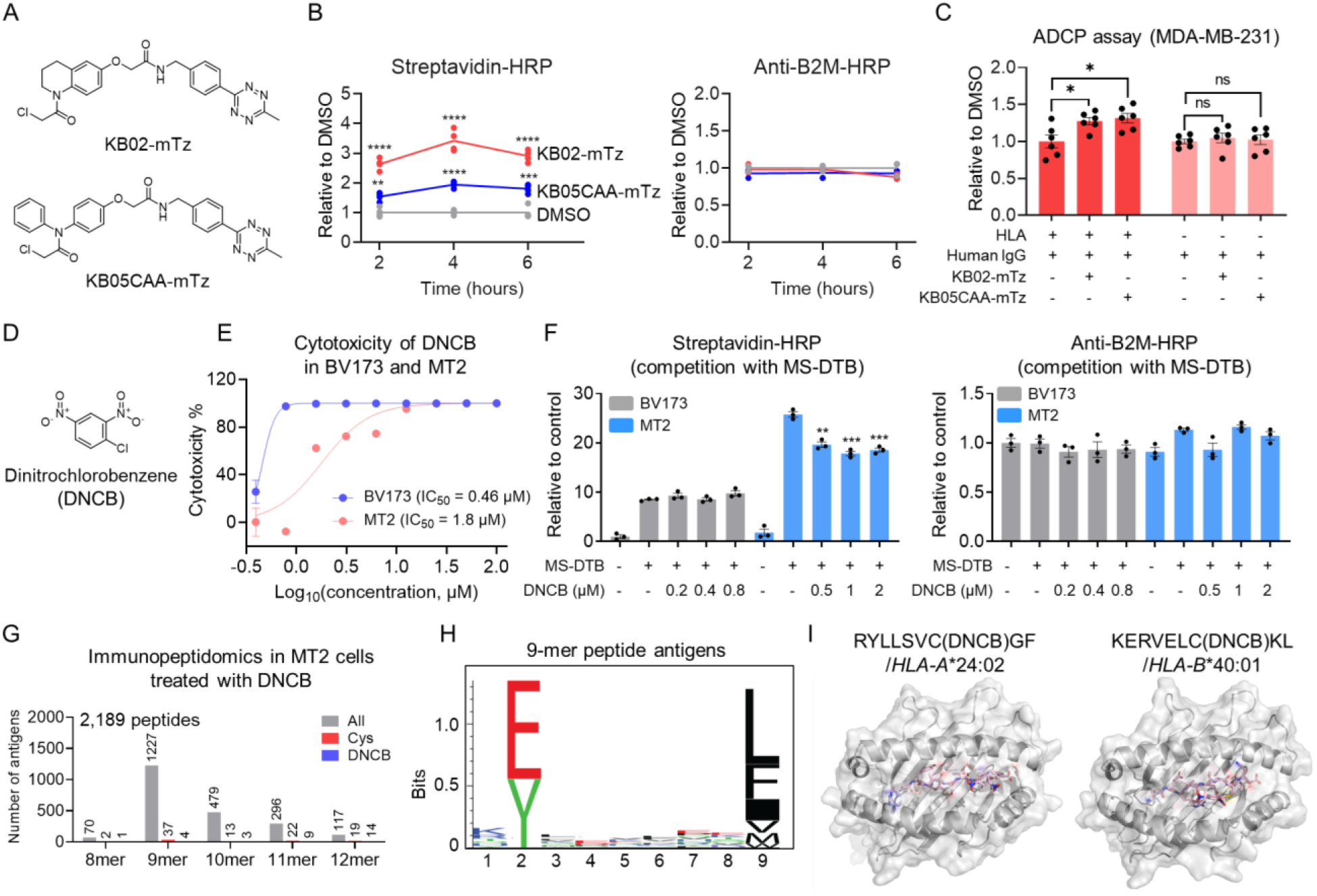
Expansion of covalent neoantigen formation using additional covalent compounds. **A**. Structures of KB02-mTz and KB05CAA-mTz. **B**. ELISA assay measuring KB02-mTz-and KB05CAA-mTz-modified peptides and B2M levels within the pMHC-I complex. Data are presented as mean ± SEM (n = 4 independent replicates for streptavidin-HRP; n = 3 for anti-B2M-HRP). The statistical significance was assessed using unpaired two-tailed Student’s t-tests. **C**. ADCP assay measuring luciferase activation in Jurkat-Luc NFAT-CD32 cells following co-culture with TCO-IgG and KB02-mTz- or KB05CAA-mTz-pretreated MDA-MB-231 *HLA* wildtype and knockout cells. Data are presented as mean ± SEM (n = 6 independent replicates). **D**. Structure of DNCB. **E**. Cytotoxicity assessment of DNCB in MT2 and BV173 cells after 72 hours of treatment. Data are presented as mean ± SEM (n = 3 independent replicates). **F**. ELISA assay measuring MSD-DTB-modified antigens within the pMHC-I complex in cells pretreated with DNCB for 24 hours. Data are presented as mean ± SEM (n = 3 independent replicates). The statistical significance was assessed using unpaired two-tailed Student’s t-tests. **G**. Distribution of 8-12-mer MHC-I-bound peptides. The result is representative of two independent experiments (n = 2). **H**. Motif analysis of all 9-mer MHC-I-bound peptide antigens. **I**. Modeling studies suggest that the DNCB-modified NPIPB13_613-621_ and SEMA3D_111-119_ peptides fit within the binding grooves of *HLA-A*\*24:02 and *HLA-B*\*40:01, respectively.

Finally, we sought to investigate 2,4-Dinitrochlorobenzene (DNCB) (**Figure 3D**), a potent sensitizer known to cause allergic contact dermatitis and affect immunocompetent individuals worldwide^19^. DNCB is an electrophilic compound, commonly produced as an intermediate in chemical manufacturing, that primarily modifies protein cysteine residues^20^. However, the link between DNCB’s cysteine reactivity and the allergic reactions it induces remains poorly understood. Given the potential widespread occurrence of covalent neoantigens, we wondered whether DNCB can modify intracellular proteins, leading to the formation of covalent neoantigens and ultimately triggering immune responses.

To this end, we first assessed the cytotoxicity of DNCB in two cell lines, MT2 and BV173, both of which express abundant pMHC-I complexes^11^. This guided the selection of appropriate concentrations to investigate covalent neoantigen formation without inducing cell death. Based on the IC_50_ values of DNCB (**Figure 3E**), we then sought to measure covalent neoantigen formation. Since DNCB lacks a bioorthogonal group, we applied a competition-based method to quantify DNCB-modified antigens. Cells were treated with maleimide-sulfonate-DTB (MS-DTB), a cell-impermeable cysteine-reactive probe previously used to directly label cysteine-containing antigens in pMHC-I complexes^11^, generating a baseline signal (**Figure 3F**). Next, cells were pretreated with DNCB for 24 hours, followed by MS-DTB labeling for 30 minutes to detect any loss of MS-DTB-modified antigen signals, indicative of DNCB modification blocking MS-DTB binding. We observed a significant decrease in signal across all DNCB concentrations (0.5, 1, and 2 µM) in MT2 cells, but not in BV173 cells (**Figure 3F**). A possible explanation for the lack of competition in BV173 cells is their lower baseline signal compared to MT2 cells, which may obscure detection of competition effects. Additionally, lower DNCB concentrations were used in BV173 cells due to their increased sensitivity (lower IC_50_ value), which may have been insufficient to produce detectable DNCB-modified antigens.

We then employed immunopeptidomics to directly identify DNCB-modified cysteine-containing peptide antigens within the MHC-I immunopeptidome. 2,189 unique 8-12-mer peptides were identified (**Figure 3G** and **Table S2**), with motif analysis of 9-mer peptides consistent with the binding preferences of MT2 *HLA* alleles (*HLA-A*\*24:02 and *HLA-B*\*40:01)^11^ (**Figure 3H**). Among four DNCB-modified 9-mer peptides, NPIPB13_613-621_ (RYLLSVCGF) and SEMA3D_111-119_ (KERVELCKL) exhibit high predicted MHC-I binding affinities in their unmodified forms (NPIPB13_613-621_ rank score for *HLA-A*\*24:02: 0.11; SEMA3D_111-119_ rank score for *HLA-B*\*40:01: 0.45). Modeling of DNCB-modified versions of these peptides showed retention of strong binding affinities for MHC-I proteins encoded by the corresponding *HLA* alleles (**Figure 3I**). Together, these data suggest a potentially widespread presence of DNCB-modified antigens within the MHC-I immunopeptidome and provide valuable targets for future investigation into the mechanisms underlying DNCB-induced immune responses.

## Discussion

This study reveals the widespread formation of covalent neoantigens across diverse cell types following treatment with cysteine-reactive electrophilic compounds. Moreover, introducing a bioorthogonal tag onto these compounds enables cells to present tagged antigens, allowing rapid and selective conjugation to immune cell engagers bearing complementary bioorthogonal groups. This strategy may offer new opportunities to develop novel modalities for immune cell recruitment and activation. Additionally, we show that the skin sensitizer DNCB facilitates the formation of covalent neoantigens, providing a potential avenue to investigate the mechanisms underlying DNCB-induced immune responses.

From a therapeutic perspective, targeting proteins for the generation of covalent neoantigens has significant potential to transform current immunotherapy strategies. A key advantage of this approach is its ability to modify proteins regardless of their function or mutational status. By chemically modifying proteins, including those that are non-essential and non-mutant, cells can present covalent neoantigens with therapeutic potential. Two recent studies demonstrated that covalent neoantigens can be harnessed to generate antibodies that recognize them, which can then be incorporated into BiTEs to recruit CTLs for the targeted elimination of cancer cells^3, 4^. While this innovative strategy opens a new avenue for targeted immunotherapy, the development of antibodies against each covalent neoantigen remains time-consuming and labor-intensive. Our work introduces a complementary approach that may enable a generalizable strategy to promote immune recognition of cancer cells presenting covalent neoantigens. By incorporating a bioorthogonal chemical group into protein-targeting inhibitors, these probes not only retain target engagement but also interfere with antigen processing, enabling the presentation of bioorthogonal covalent neoantigens on the cell surface. This bioorthogonal handle can be further exploited to develop universal modalities that engage diverse immune cells for therapeutic applications. In this study, we demonstrated the feasibility of recruiting Fc receptor-expressing immune cells. Future designs could incorporate CD3-binding single-chain variable fragments (scFvs) to construct universal T cell engagers capable of recruiting CTLs to eliminate target cells treated with a broad range of probes that generate bioorthogonal covalent neoantigens.

Moving forward, we are interested in exploring a wide range of well-characterized, target-specific inhibitors and assessing their potential to be converted into bioorthogonal probes capable of generating covalent neoantigens. These probes could then be paired with immune cell engagers for targeted cell elimination. We also hypothesize that this mechanism may extend to other covalent compounds that target non-cysteine nucleophilic residues, such as lysine, serine, histidine, and tyrosine. Investigating chemical probes that engage these residues could further expand the repertoire of chemically induced covalent neoantigens—not only enhancing our understanding of this emerging class of antigens but also unlocking new therapeutic opportunities.

In this study, we investigated DNCB, a skin sensitizer, and found that it can lead to the formation of covalent neoantigens. Given that many allergenic chemicals and drugs, or their reactive metabolites, are covalent molecules capable of forming protein adducts^21^, it is possible that such modifications generate covalent neoantigens presented on MHC-I, thereby triggering immune responses^22^. However, this long-standing hypothesis has not been systematically investigated due to technological limitations, creating a bottleneck in our understanding of chemical and drug-induced allergies. Our platform provides a way to address this barrier and advance our understanding of the immunological mechanisms underlying these allergic responses, an essential step toward developing future strategies for their prevention or mitigation.

## Acknowledgement

We gratefully acknowledge the support of the Ono Pharma Foundation (X.Z.) and Ryan Family Research Acceleration Fund Award (X.Z.).

## Author Contributions Statement

C.Zhou and C. Zhang designed and conducted biochemical and cellular experiments; C.Zhou, W.H., and X.J. synthesized and characterized the compounds; H.W. conducted ELISA assays; C.Zhang conducted the modeling study; C.Zhou and C.Zhang conducted immunopeptidomics experiments; C.Zhou, C.Zhang, L.M.M., and M.S. performed data analysis and identified candidate targets for the immunopeotidomics studies; C.Zhang and X.Z. supervised the project and, with contributions from all authors, wrote the manuscript.

## Data Availability

The mass spectrometry proteomics data were deposited to the ProteomeXchange Consortium through the PRIDE^23^ partner repository with the dataset identifier PXD065352. The data supporting the findings of this study are available within the article and Supplementary Information. Source data are provided with this paper.

## Supplementary Figures

**Figure S1.**
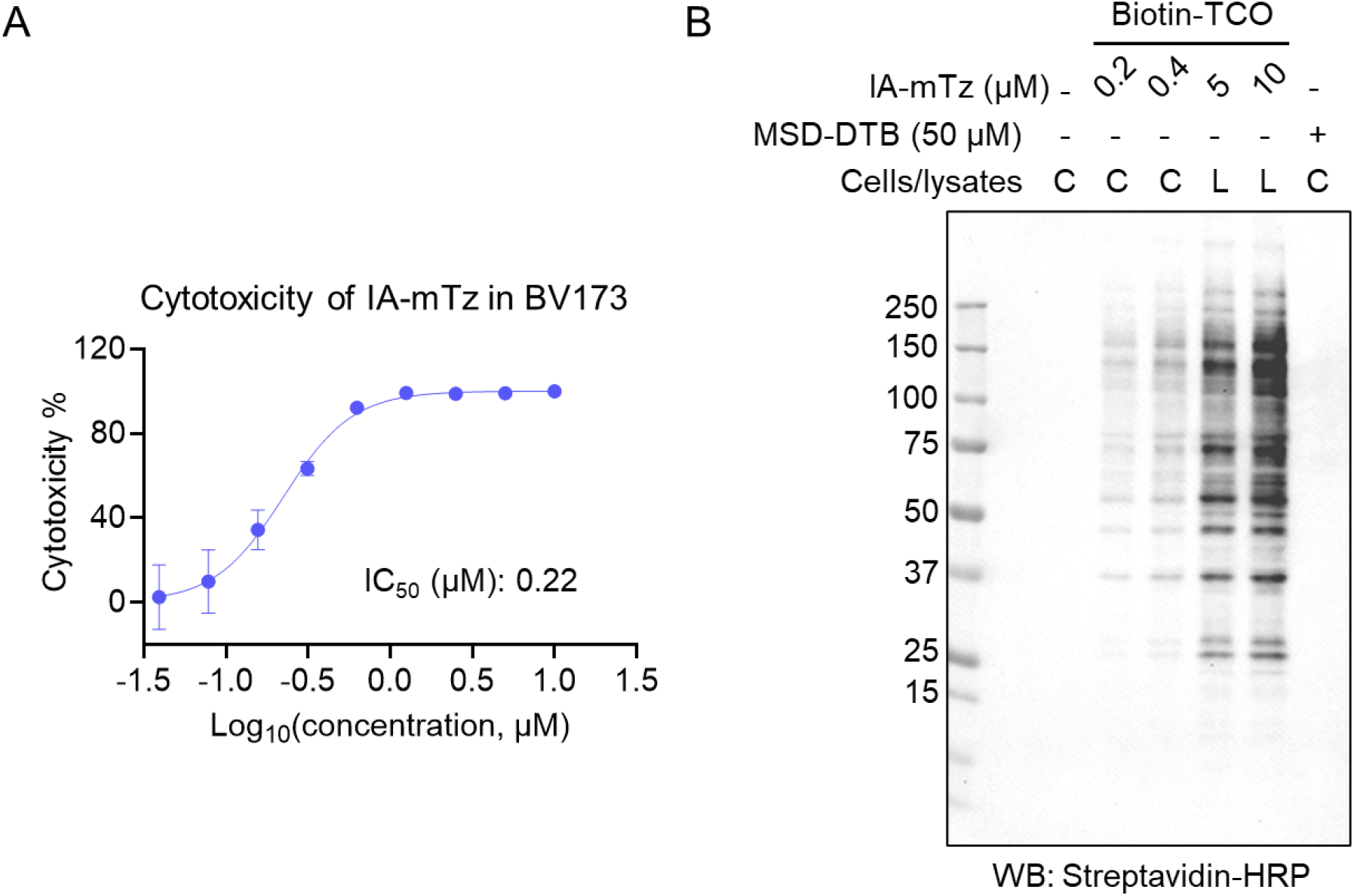
Development of a cell-permeable probe to identify covalent neoantigens. **A**. Cytotoxicity assessment of IA-mTz in BV173 cells after 72 hours of treatment. Data are presented as mean ± SEM (n = 3 independent replicates). **B**. Western blot analysis revealed that IA-mTz effectively labeled proteins in live cells. Maleimide-sulfonate-dibenzocyclooctyne-DTB (MSD-DTB), a previously reported cell-impermeable and cysteine-reactive probe^1^, serves as a control and shows no detectable labeling in live cells.

**Figure S2.**
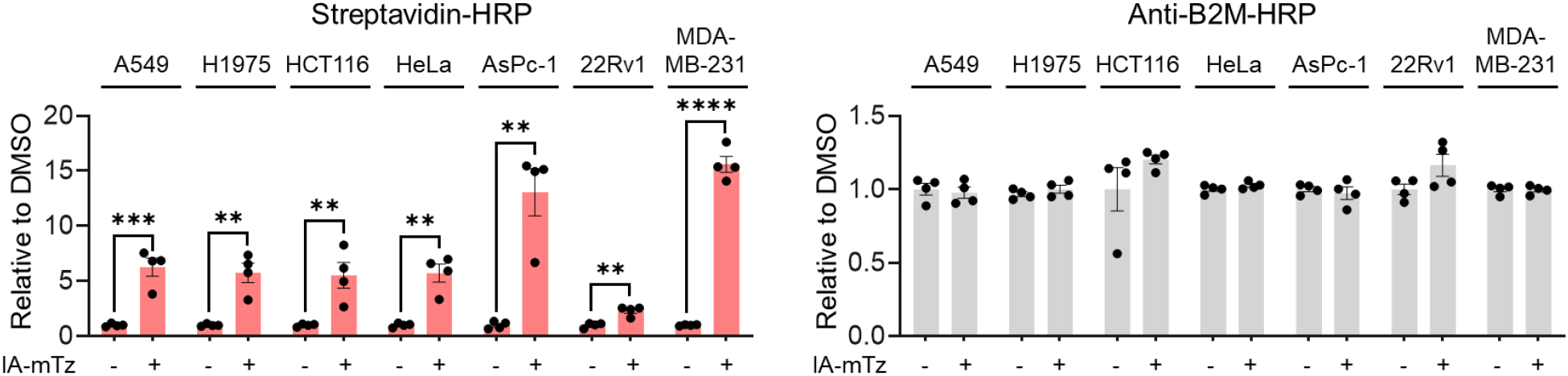
ELISA assay measuring IA-mTz-modified peptides and B2M levels within the pMHC-I complex in seven cell lines. Data are presented as mean ± SEM (n = 4 independent replicates). The statistical significance was assessed using unpaired two-tailed Student’s t-tests.

**Figure S3.**
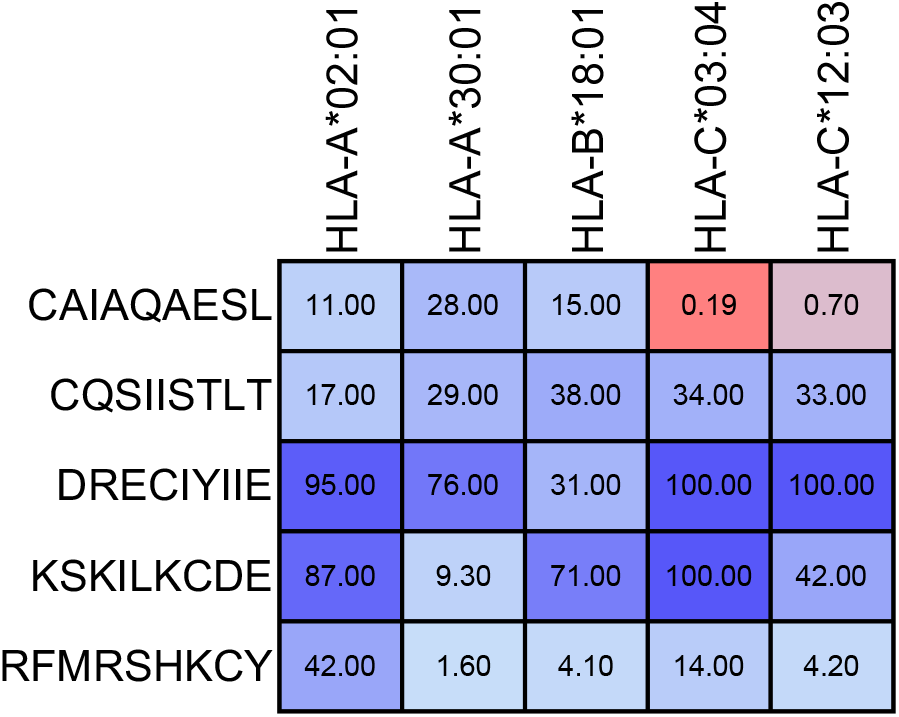
Predicted MHC-I binding scores for the five IA-mTz-modified 9-mer antigens across HLA alleles expressed in BV173 cells (*HLA-B*\*15:10 not available in IEDB).

**Figure S4.**
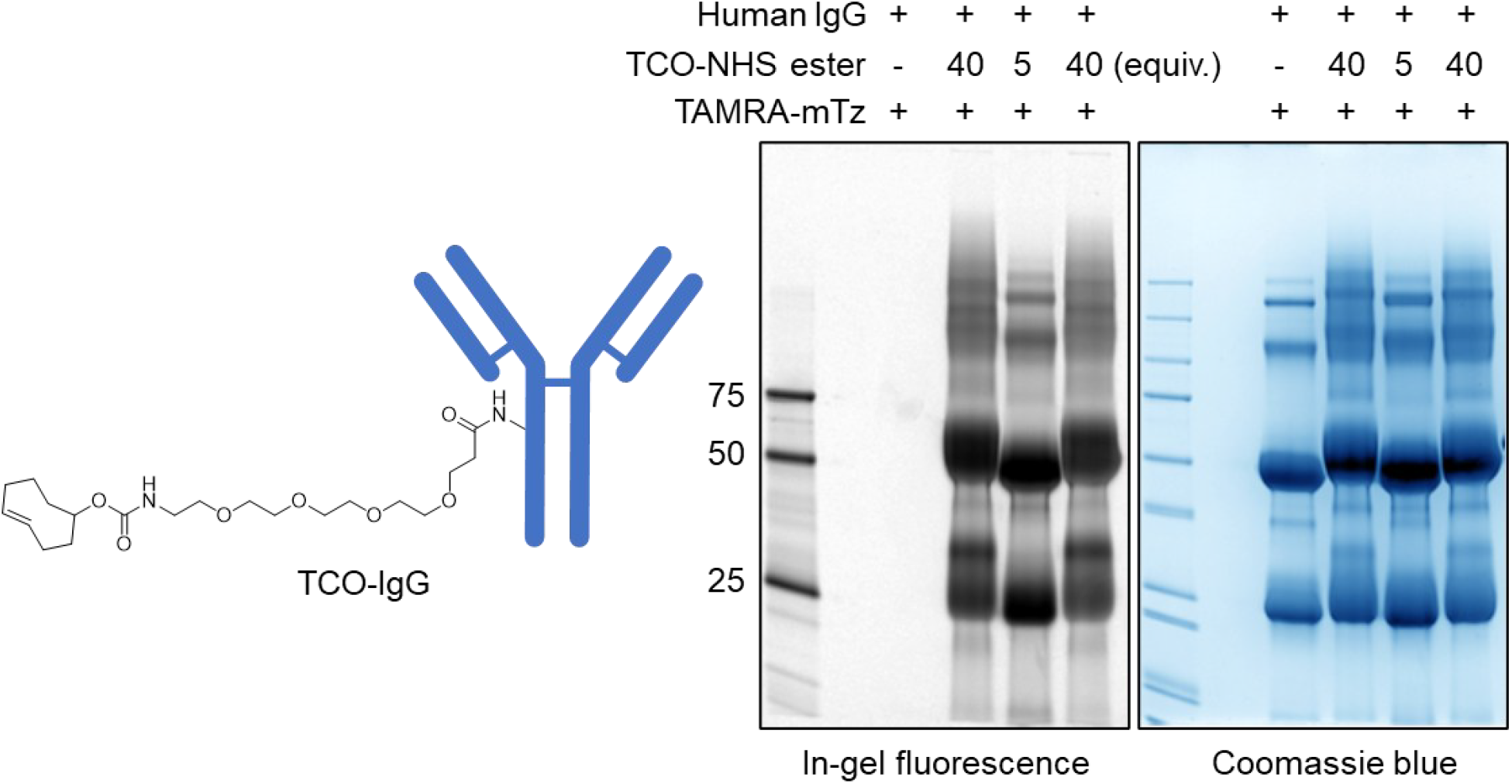
In-gel fluorescence analysis confirmed the conjugation of TCO to IgG.

**Figure S5.**
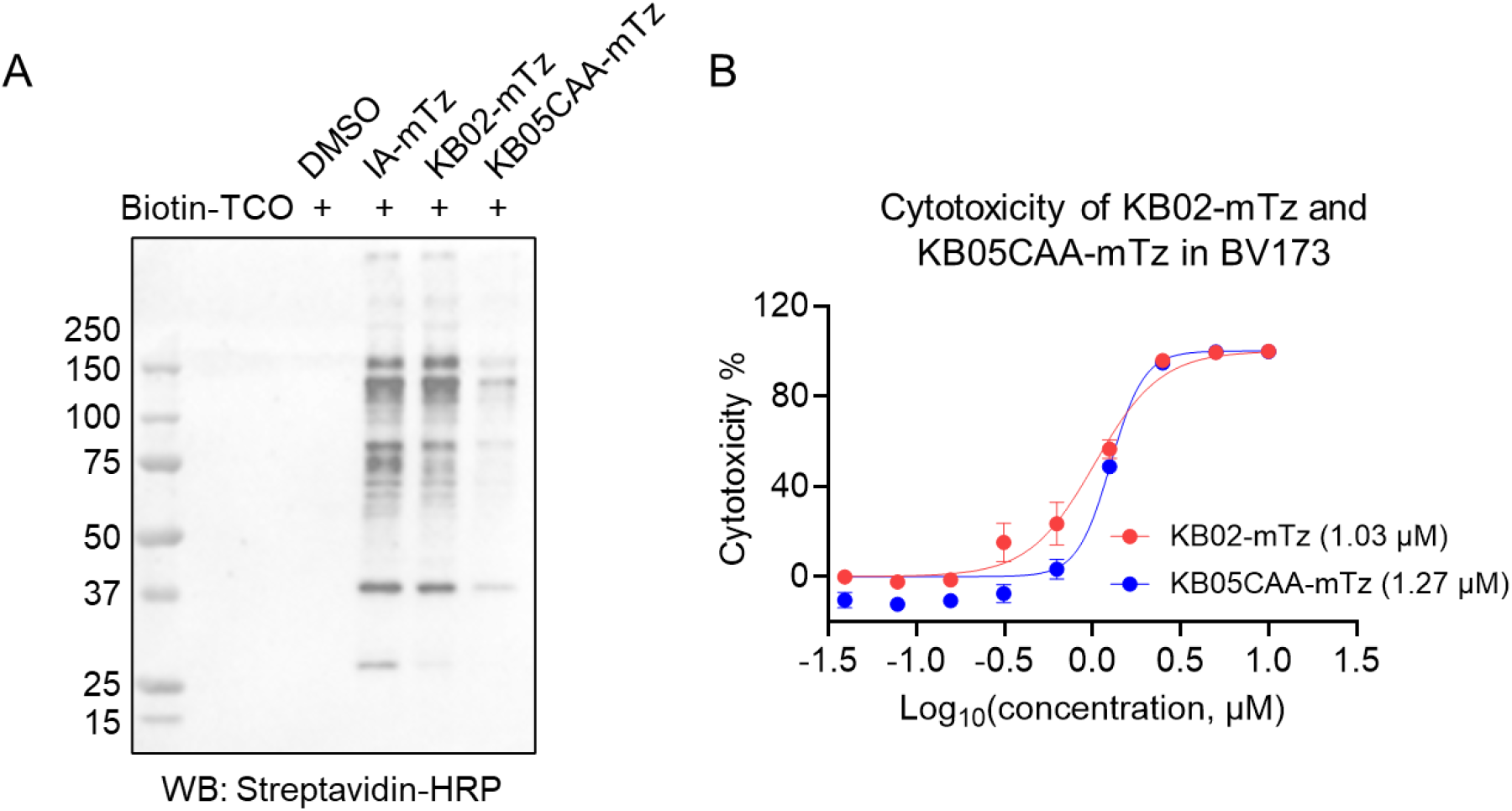
Expanding covalent neoantigens with KB02-mTz and KB05CAA-mTz. **A**. Western blot analysis revealed that IA-mTz effectively labeled proteins in live cells. **B**. Cytotoxicity assessment of KB02-mTz and KB05CAA-mTz in BV173 cells after 72 hours of treatment. Data are presented as mean ± SEM (n = 3 independent replicates).

