## Supplementary Information for "Expanding the repertoire of chemically induced covalent neoantigens"

### Materials and Methods

#### Reagents

Streptavidin-HRP (Cat. #3999) was purchased from Cell Signaling Technology. InVivoMAb anti-human MHC Class I (HLA-A, HLA-B, HLA-C) antibody (clone W6/32, Cat. #BE0079) used for ELISA assays was purchased from Bio X Cell. Ultra-LEAF™ anti-human HLA-A, B, C antibody (clone W6/32, Cat. #311448) used for immunopeptidomics was purchased from BioLegend. Human IgG isotype (Cat. #31154), Pierce™ enzyme-linked chemiluminescence (ECL) western blotting substrate (Cat. #32106), SuperSignal™ West Pico PLUS Chemiluminescent Substrate (Cat. # 34577), QuantaBlu™ fluorogenic peroxidase substrate kit (Cat. #15169), and beta-2-microglobulin HRP (clone B2M-01, Cat. #MA1-19679) were purchased from Thermo Scientific. HLA-C*03:04&B2M (Cat. #HLM-H82Er) and HLA-A*02:01&B2M (Cat. #HLM-H52H5) recombinant proteins were purchased from ACROBiosystems. TCO-PEG4-NHS (Cat. #BP-22418), IA-mTz (Cat. #BP-41387), Biotin-TCO (Cat. #BP-23847) were purchased from BroadPharm.

#### Cell Lines

A549, H1975, HCT116, HeLa, AsPc-1, 22Rv-1, and MDA-MB-231 cells were obtained from ATCC. BV173 cells were obtained from CLS Cell Lines Service. MT2 cells were obtained from Thermo Scientific. Jurkat-Lucia NFAT-CD32 cells were obtained from InvivoGen. A549, HeLa, MDA-MB-231 cells were cultured in Dulbecco's Modified Eagle Medium (DMEM, Corning) with 10% (v/v) fetal bovine serum (FBS, Omega Scientific) and L-glutamine (2 mM, Gibco). BV173, and H1975, AsPc-1, and 22Rv-1 cells were cultured in RPMI 1640 (Corning) with 10% (v/v) FBS (Omega Scientific) and L-glutamine (2 mM, Gibco). HCT116 cells were cultured in McCoy's 5A Medium (Corning) with 10% (v/v) fetal bovine serum (FBS, Omega Scientific) and L-glutamine (2 mM, Gibco). Jurkat-Lucia NFAT-CD32 cells were cultured in Iscove's Modification of DMEM (Corning) with 10% (v/v) FBS (Omega Scientific) and L-glutamine (2 mM, Gibco). All the cell lines were tested negative for mycoplasma contamination using Universal Mycoplasma Detection Kit (ATCC, cat# 30-1012K).

#### Generation of CRISPR-Cas9-mediated *HLA* knockout cells

MDA-MB-231 cells with CRISPR-Cas9-mediated knockout of *HLA-A,B,C* were generated by electroporation of a Cas9-sgRNA ribonucleoprotein (RNP) complex using the 4D-Nucleofector system (Lonza Bioscience). A mixture of three sgRNAs targeting *HLA* genes was used for electroporation: sgRNA#1 (CGGCTACTACAACCAGAGCG), sgRNA#2 (AGATCACACTGACCTGGCAG), and sgRNA#3 (AGGTCAGTGTGATCTCCGCA). The *HLA* knockout was validated in a previously published study[1].

**Cell Lysis and Western Blot**

Cells were pretreated at 37 °C with either 200 nM or 400 nM IA-mTz, 1 μM KB02-mTz, 1 μM KB05CAA-mTz for 2 hours, or 50 μM MSD-DTB for 30 minutes, then washed twice with Dulbecco's Phosphate-Buffered Saline (DPBS) and harvested. Cells were lysed in DPBS supplemented with cOmplete protease inhibitor cocktail (Roche). The cell suspension was subjected to sonication for 5 cycles at 40% power, with 4 pulses per cycle. For in-lysate labeling, following sonication, the lysate was incubated with 5 or 10 μM IA-mTz for 1 hour at room temperature. Subsequently, 20 μM biotin-TCO was added and the mixture was incubated for an additional hour at room temperature. The reaction mixture was then centrifuged at 16,000 × g for 10 minutes at 4 °C to collect the supernatant. Protein concentration in the supernatant was determined using the DC Protein Assay (Bio-Rad). The protein lysate was mixed with Laemmli sample buffer (Bio-Rad) and heated at 95 °C for 5 minutes. Proteins were separated on 4-20% Novex Tris-Glycine mini gels (Invitrogen), then transferred onto a 0.2 μm polyvinylidene fluoride (PVDF) membrane (Bio-Rad). The membrane was blocked with 5% non-fat milk in Tris-buffered saline with 0.1% Tween 20 (TBST; 20 mM Tris-HCl, pH 7.6, 150 mM NaCl) for 1 hour at room temperature. Streptavidin-HRP was diluted 1:1000 in 5% non-fat milk/TBST and incubated with the membrane for 1 hour at room temperature. After three washes with TBST, the chemiluminescent signal was developed using ECL Western blotting detection reagent or SuperSignal West Pico PLUS Chemiluminescent Substrate and imaged with a ChemiDoc MP system (Bio-Rad).

**Cell Viability Assay**

Cells were seeded in a 96-well clear-bottom white plate (Corning) at a density of 10,000 cells per well in 100 μL of RPMI medium and incubated for 24 hours. The cells were then treated with varying concentrations of compounds diluted in 100 µL of RPMI medium for an additional 72 hours. After treatment, 50 µL of CellTiter-Glo reagent (Promega) was added to each well and incubated for 10 minutes at room temperature. Luminescence was measured using a CLARIOstar Plus microplate reader (BMG Labtech).

**ELISA Assay for Detecting Tetrazine Modification**

Cells were treated with IA-mTz, KB02-mTz, or KB05CAA-mTz at 250 nM for BV173 cells, 2 µM for H1975 and 22Rv1 cells, or 5 µM for A549, HCT116, HeLa, AsPC-1, and MDA-MB-231 cells, then washed twice with DPBS and harvested. Nunc MaxiSorp 384-well black plates were coated overnight with 50 µL of anti-heavy chain antibody W6/32 at 5 µg/mL in PBS. After coating, plates were washed twice with 100 µL PBS and blocked with 120 µL of 3% bovine serum albumin (BSA) in PBS for 1 hour at room temperature. Plates were then washed three times with 100 µL of 0.05% Tween-20 in PBS (PBST). Treated cells were lysed in NP-40 lysis buffer containing protease inhibitors. Protein concentration was adjusted to 1 mg/mL, and 50 µL of total lysate or MHC-I refolding mixture was added per well. Plates were incubated at room temperature for 1 hour, followed by three washes with 100 µL of 1% BSA in PBS. Next, 50 µL of either 1 µM biotin-TCO in 1% BSA PBS or 1% BSA PBS (control) was added to each well. Plates were incubated with shaking at room temperature for 1 hour, then washed twice with PBST and three times with PBS (100 µL per wash). Next, 50 µL of either 1 µg/mL anti-beta-2-microglobulin-HRP conjugate or Streptavidin-HRP (1:1000 dilution) in 1% BSA PBS was added to each well. Plates were incubated with shaking for 1 hour at room temperature. After incubation, plates were washed three times with PBST and three times with PBS (100 µL each). Finally, 50 µL of the HRP substrate QuantaBlu was added, and fluorescence was measured using a CLARIOstar Plus microplate reader (BMG Labtech).

**ELISA Assay for Detecting DNCB Modification**

Nunc MaxiSorp 384-well black plates were coated overnight with 50 µL of anti-heavy chain antibody W6/32 at 5 µg/mL in PBS. After coating, plates were washed twice with 100 µL PBS and blocked with 120 µL of 3% bovine serum albumin (BSA) in PBS for 1 hour at room temperature. Plates were then washed three times with 100 µL of 0.05% Tween-20 in PBS (PBST). BV173 and MT2 cells were treated with DNCB (0.2, 0.4, 0.8 µM for BV173; 0.5, 1.0, and 2.0 µM for MT2) for 24 hours, followed by treatment with 50 µM

MS-DTB. After washing, cells were harvested and lysed in NP-40 lysis buffer containing protease inhibitors. Protein concentration was adjusted to 1 mg/mL, and 50 µL of total lysate was added to each well. Plates were incubated at room temperature for 1 hour, then washed three times with 100 µL of 1% BSA in PBS. Next, 50 µL of either 1 µg/mL anti-beta-2-microglobulin-HRP conjugate or Streptavidin-HRP (1:1000 dilution) in 1% BSA PBS was added to each well. Plates were incubated with shaking at room temperature for 1 hour, followed by three washes with PBST and three washes with PBS (100 µL each). Finally, 50 µL of HRP substrate QuantaBlu was added, and fluorescence was measured using a CLARIOstar Plus microplate reader (BMG Labtech).

**MSD Blocking Experiment**

BV173 cells (2 × $10^6$ cells/mL) were pretreated with DMSO or 50 µM MSD for 30 minutes. Subsequently, 25 µM MSD-DTB or 250 nM IA-mTz were added and incubated for 2, 4, or 6 hours. Every 2 hours, an additional 25 µM MSD was added to the culture to maintain continuous blocking. Cells were then harvested for ELISA assays as described above.

**MHC-I Refolding Assay**

HLA-C*03:04&B2M and HLA-A*02:01&B2M monomer proteins were dissolved in denaturation buffer (8 M urea, 20 mM Tris-HCl, pH 8.0, and 10 mM DTT) at a final concentration of 0.5 mg/mL and incubated overnight at 4 °C. The denatured HLA&B2M monomers were then diluted 100-fold into refolding buffer (100 mM Tris-HCl, pH 8.0; 400 mM L-arginine-HCl; 2 mM EDTA; 5 mM reduced glutathione; and 0.5 mM oxidized glutathione) and incubated at 4 °C for 30 minutes. Subsequently, peptide antigen was added to the refolding mixture at a molar ratio of HLA to peptide of 1:10. The mixture was incubated at 4 °C for 24 hours before being subjected to ELISA assay.

**Immunopeptidomics**

8 × $10^8$ BV173 or MT2 cells (2 × $10^6$ cells/mL) were treated with either 250 nM IA-mTz continuously for 1 week or 2 µM DNCB for 24 hours. Cells were harvested and lysed in 8 mL of lysis buffer (0.5% NP-40, 50 mM Tris-HCl pH 8.0, 150 mM NaCl, 1 mM EDTA, and protease inhibitor cocktail) by rotating at 4 °C for 30 minutes. Following centrifugation at 18,000 × g for 10 minutes, the supernatant was collected for enrichment using an anti-

MHC antibody (W6/32, BioLegend) conjugated to Affi-Gel 10 matrix (Bio-Rad; 2 mg of antibody per 100 μL of slurry per sample). Enrichment was performed by rotating the lysate with the resin at 4 °C for 4 hours, followed by transfer to a Bio-Spin column (Bio-Rad). Columns were sequentially washed with 3 × 1 mL of lysis buffer, wash buffer 1 (50 mM Tris-HCl pH 8.0, 150 mM NaCl), wash buffer 2 (50 mM Tris-HCl pH 8.0, 400 mM NaCl), and wash buffer 3 (50 mM Tris-HCl pH 8.0). MHC-bound peptides were eluted with 1 mL of 1% trifluoroacetic acid in water. Peptide samples were desalted using Sep-Pak C18 cartridges (Waters), dried by SpeedVac, and analyzed on an Orbitrap Eclipse Tribrid mass spectrometer coupled with a Vanquish Neo UHPLC system.

Peptides were loaded onto an EASY-Spray HPLC column (C18, 2 μm particle size, 75 μm inner diameter, 150 mm length) and eluted at a flow rate of 0.25 μL/min using the following gradient: 5% buffer B (80% acetonitrile with 0.1% formic acid) in buffer A (water with 0.1% formic acid) from 0 to 15 minutes; increasing to 45% buffer B from 15 to 155 minutes; and then from 45% to 100% buffer B from 155 to 180 minutes. The nano-LC electrospray ionization source was operated at 1.5 kV. The analysis began with an MS1 scan in the Orbitrap (resolution 120,000; m/z range 375–1600; RF lens at 30%; standard AGC target; automatic maximum injection time). Precursor ions were isolated in MS2 using the quadrupole with a 0.7 m/z isolation window and fragmented via higher-energy collisional dissociation (HCD) in the ion trap (collision energy 27%; standard AGC target; maximum injection time 35 ms). RAW data were acquired using Xcalibur software (version 4.5.445.18) and analyzed with Proteome Discoverer version 2.5. Peptide motif analysis was conducted using the Seq2Logo method.

**Generation of TCO-IgG**

Human IgG isotype (Thermo Scientific) was diluted in PBS (pH 8.0) to a final concentration of 25 μM. TCO-PEG4-NHS was then added to a final concentration of 1 mM, and the reaction mixture was incubated overnight at room temperature. Unreacted TCO-PEG4-NHS was removed using Amicon Ultra Centrifugal Filters (30 kDa MWCO, MilliporeSigma), and the buffer was exchanged to PBS (pH 7.4). For in-gel fluorescence analysis, 10 μg of modified or unmodified Human IgG isotype was incubated with TAMRA-mTz at room temperature for 1 hour. Laemmli sample buffer was then added, and the mixture was heated at 95 °C for 10 minutes. Samples were resolved by 4-20% Novex

Tris-Glycine mini gels, and fluorescence signals were recorded using the ChemiDoc MP system.

**Antibody-Dependent Cellular Phagocytosis Assay**

200 µL of MDA-MB-231 cells or H1975 cells at $5 \times 10^4$ cells per well were seeded in a 96-well plate and incubated overnight at 37 °C. The following day, cells were treated with either DMSO or 5 µM of IA-mTz, KB02-mTz, or KB05CAA-mTz and incubated overnight. After treatment, cells were washed three times with fresh media, followed by the addition of 100 µL of TCO-IgG (100 µg/mL) per well and incubated for 1 hour at 37 °C. Subsequently, 100 µL of Jurkat-Luc NFAT-CD32 cells ($8 \times 10^5$ cells per well) were added to each well and incubated overnight at 37 °C. After incubation, 20 µL of culture supernatant was transferred to a 96-well white plate and mixed with 50 µL of QUANTI-Luc 4 Lucia reagent (InvivoGen) per well. Luciferase activity was measured using a CLARIOstar Plus microplate reader.

**Modeling Study**

The RS3$^{97\text{-}105}$-HLA-C*03:04 complex, along with the structures of HLA-A*24:02 and HLA-B*40:01 bound to two different peptides, NPIPB13$^{613\text{-}621}$ (RYLLSVCGF) and SEMA3D$^{111\text{-}119}$ (KERVELCKL), were predicted using AlphaFold 3. Structural preprocessing, including hydrogen-bond optimization, energy minimization, and removal of water molecules, was performed using the Protein Preparation Wizard in Maestro 13.4 (Schrödinger). Ligands were prepared with the LigPrep module employing the OPLS4 force field. Covalent docking was carried out using the CovDock module, with the nucleophilic substitution reaction type selected and default parameters applied for other settings. The final pose was selected based on low-energy conformation and favorable hydrogen-bonding and cation-pi interaction geometries. Figures were generated using PyMOL.

**Statistical Analysis**

Quantitative data were presented as scatter plots, with means indicated and error bars representing the standard error of the mean (SEM). Differences between two groups were evaluated using an unpaired, two-tailed Student's t-test. Statistical significance was defined as P < 0.05. Significance levels were indicated as follows: *$P$ < 0.05, **$P$ < 0.01, ***$P$ < 0.001, ****$P$ < 0.0001, and *ns* (not significant).

**Synthesis of MSD, MSD-DTB, and MS-DTB**

MSD, MSD-DTB, and MS-DTB were synthesized as previously described[1].

**Synthesis of KB02-mTz**

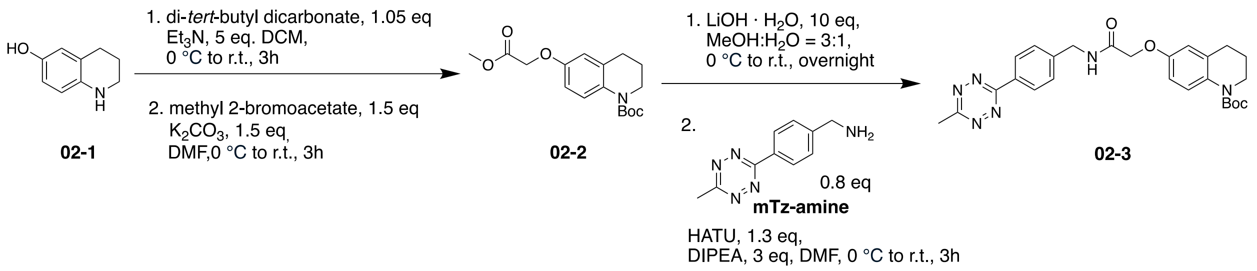


**Step 1:** 1,2,3,4-tetrahydroquinolin-6-ol, **02-1** (1 g, 6.7 mmol, 1.0 eq) and $Et_3N$ (4.68 mL, 33.5 mmol, 5 eq) were dissolved in DCM (10 mL) and stirred on ice for 30 minutes. Then di-tert-butyl decarbonate (1.6 mL, 7.035 mmol, 1.05 eq) was added and stirred 3 hours at room temperature. Upon completion, the reaction was concentrated, and the crude was extracted with EtOAc, then concentrated and purified by flash chromatography. Next, Boc protected **02-1** (1 g, 4 mmol, 1 eq) and $K_2CO_3$ (829.2 mg, 6 mmol, 1.5 eq) was dissolved in DMF (10 mL) and stirred on ice for 30 minutes. Then methyl 2-bromoacetate (1.56 mL, 6 mmol, 1.5 eq) was added stirred 3 hours at room temperature. Upon completion, the reaction was concentrated, and the crude was extracted with EtOAc. The crude mixture was concentrated and purified by flash chromatography to yield **02-2** as with powder (969 mg, 3 mmol, 45% over two steps).

**$^1$H NMR** (400 MHz, $CDCl_3$) δ 7.55 (d, $J$ = 8.8 Hz, 1H), 6.70 (dd, $J$ = 9.0, 3.0 Hz, 1H), 6.63 (d, $J$ = 3.0 Hz, 1H), 4.59 (s, 2H), 3.80 (s, 3H), 3.71 – 3.64 (m, 2H), 2.73 (t, $J$ = 6.6 Hz, 2H), 1.90 (p, $J$ = 6.4 Hz, 2H), 1.51 (s, 9H).

**Step 2:** **02-2** (80 mg, 0.25 mmol, 1 eq) and LiOH·$H_2O$ (105 mg, 2.5 mmol, 10 eq) were dissolved in MeOH:$H_2O$ (3:1, 2 mL) on ice and stirred at room temperature for overnight. Upon completion, the reaction was concentrated and extracted by DCM:MeOH (10:1).

The crude hydrolyzed product was concentrated and moved onto the next step without further purification. The crude hydrolyzed product, mTz-amine (50 mg, 0.21mmol, 0.8 eq), and HATU (103.8 mg, 0.27 mmol, 1.3 eq) was dissolved in DMF (2 mL) on ice. The DIPEA (60.4 µL, 0.63 mmol, 3 eq) was added and stirred at room temperature for 3 hours. Upon completion, the reaction was concentrated and extracted by EtOAc and purified by flash chromatography to yield **02-3** as red powder (74 mg, 0.15 mmol, 61% over two steps).

**$^1$H NMR** (400 MHz, $CDCl_3$) δ 8.59 – 8.53 (m, 2H), 7.59 (d, *J* = 9 Hz, 1H), 7.53 – 7.46 (m, 2H), 7.00 (d, *J* = 3.2 Hz, 1H), 6.73 (dd, *J* = 9, 3.0 Hz, 1H), 6.64 (dd, *J* = 6.0, 3.0 Hz, 1H), 4.69 – 4.60 (m, 2H), 3.72 – 3.64 (m, 2H), 3.10 (s, 2H), 2.73 (t, *J* = 6.4 Hz, 2H), 1.96 – 1.85 (m, 2H), 1.64 (s, 3H), 1.51 (s, 9H).

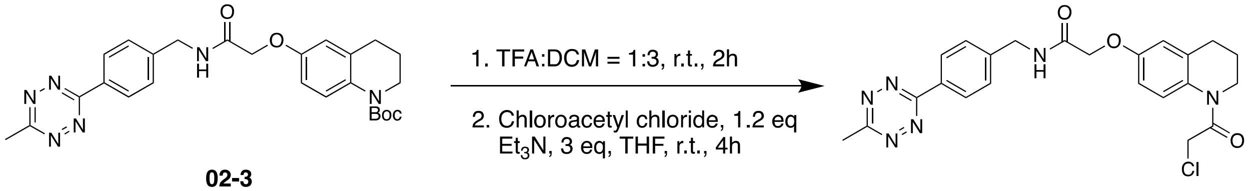


**02-3** **KB02-mTz**

**Step 3: 02-3** (50 mg, 0.1 mmol, 1 eq) was deprotected by dissolved in DCM (1 mL) and the mixture was allowed to cool to 0 °C before then slowly adding TFA (0.3 mL). The combined reaction mixture was then allowed to stir at room temperature for 2 hours. Upon completion, the reaction was concentrated under reduced pressure without further purification. Next, deprotected product and $Et_3N$ (42 µL, 0.3 mmol, 3 eq) were dissolved in THF (1.5 mL) and the mixture was allowed to cool to 0 °C before then adding chloroacetyl chloride (20 µL, 0.12 mmol, 1.2 eq) in a dropwise manner. The mixture was stirred at room temperature for 4 hours. Upon completion, the reaction was concentrated and extracted by DCM. Then the crude product was concentrated and purified by prep-TLC plate to yield **KB02-mTz** as red powder (24.7 mg, 0.053 mmol, 53% over two steps).

**$^1$H NMR** (500 MHz, $CDCl_3$) δ 8.56 – 8.51 (m, 2H), 7.49 (d, *J* = 8 Hz, 2H), 7.14 (s, 1H), 7.04 (t, *J* = 6 Hz, 1H), 6.77 (dt, *J* = 13.5, 7 Hz, 2H), 4.65 (d, *J* = 6 Hz, 2H), 4.57 (s, 2H), 4.17 (s, 2H), 3.78 (t, *J* = 6.5 Hz, 2H), 3.08 (s, 3H), 2.69 (s, 2H), 1.97 (s, 2H).

**$^{13}$C NMR** (126 MHz, $CDCl_3$) δ 168.24, 167.47, 163.93, 142.68, 131.32, 128.53 (2C), 128.45 (2C), 114.68, 112.93, 67.74, 42.81, 29.83, 28.21, 27.04, 23.74, 21.31.

**HRMS (ESI+)** *m/z* calcd for $C_{23}H_{23}ClN_6O_3Na^+$ $[M+Na]^+$: 489.1418, found 489.1446.

#### Synthesis of KB05CAA-mTz

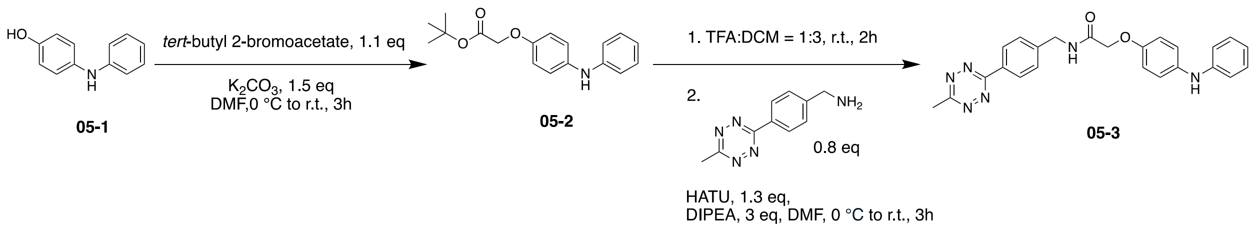


**Step1:** 4-(phenylamino)phenol, **05-1** (1 g, 5.4 mmol, 1 eq) and $K_2CO_3$ (1.12g, 8.1 mmol, 1.5 eq) were dissolved in DMF (20 mL) and stirred on ice for 30 minutes. Then tert-butyl 2-bromoacetate (1.53 mL, 5.94 mmol, 1.1 eq) was added and stirred at room temperature for 3 hours. Upon completion, the reaction was concentrated and extracted by EtOAc. The combined organic layer was then concentrated and purified by flash chromatography to yield 05-2 as brown oil (1.4g, 4.7mmol, 87%).

**$^1$H NMR** (500 MHz, DMSO-$d_6$) δ 7.86 (s, 1H), 7.20 – 7.12 (m, 2H), 7.05 – 6.98 (m, 2H), 6.92 (dq, *J* = 7, 1.5 Hz, 2H), 6.86 – 6.79 (m, 2H), 6.72 (tt, *J* = 7.5, 1.0 Hz, 1H), 4.57 (s, 2H), 1.43 (s, 9H).

**Step 2:** Tert-butyl ester in **05-3** was hydrolyzed by dissolved it in DCM:TFA = 3:1 on ice and stirred at room temperature for 2 hours. Upon completion, the reaction was concentrated under reduced pressure without further purification. Next, the hydrolyzed product (38.4 mg, 0.158 mmol, 1 eq), **mTz-amine** (30.04 mg, 0.126 mmol, 0.8 eq), and HATU (63.3 mg, 0.205 mmol, 1.3 eq) was dissolved in DMF (1 mL) on ice. The DIPEA (82.6 μL, 0.474 mmol, 3 eq) was added and stirred at room temperature for 3 hours. Upon completion, the reaction was concentrated and extracted by EtOAc and purified by flash chromatography to yield **05-3** as red oil (22.2 mg, 0.052 mmol, 33%).

**$^1$H NMR** (400 MHz, $CDCl_3$) δ 8.60 – 8.53 (m, 2H), 7.50 (d, *J* = 8.2 Hz, 2H), 7.23 (dd, *J* = 8.4, 7.2 Hz, 2H), 7.11 – 7.00 (m, 3H), 6.98 – 6.91 (m, 2H), 6.87 (dd, *J* = 8.0, 5.6 Hz, 3H), 5.55 (s, 1H), 4.67 (d, *J* = 6 Hz, 2H), 4.58 (s, 2H), 3.10 (s, 3H).

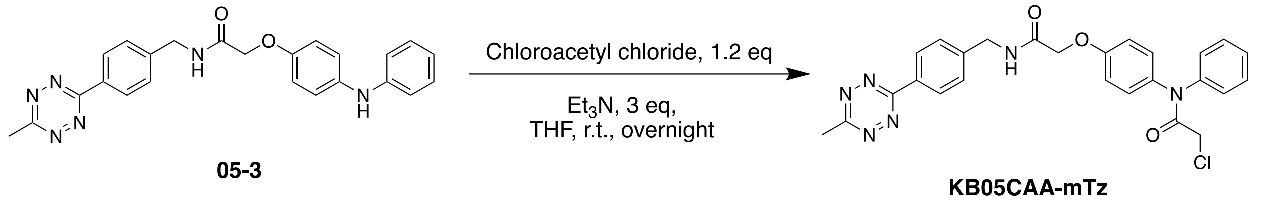

**Step 3: 05-3** (15 mg, 0.035 mmol, 1 eq), and $Et_3N$ (19.5 μL, 0.14 mmol, 3 eq) were dissolved in THF (0.5 mL) and the mixture was allowed to cool to 0 °C before then adding chloroacetyl chloride (4 μL, 0.053 mmol, 1.5 eq) in a dropwise manner. Then mixture was stirred at room temperature overnight. Upon completion, the reaction was concentrated and extracted by DCM. Then the crude product was concentrated and purified by Prep-TLC plate to yield **KB05CAA-mTz** as red oil (6.7 mg, 0.013 mmol, 37%).

**$^1$H NMR** (500 MHz, $CDCl_3$) δ 8.53 (d, J = 8.0 Hz, 2H), 7.48 (d, J = 8.0 Hz, 9H), 7.08 – 6.83 (m, 3H), 4.65 (d, J = 6.0 Hz, 2H), 4.57 (s, 2H), 4.00 (s, 2H), 3.09 (s, 3H).

**$^{13}$C NMR** (126 MHz, $CDCl_3$) δ 168.08, 167.44, 166.41, 163.92, 142.63, 141.84, 131.29, 130.27, 129.24, 128.50, 128.43, 127.74, 125.92, 116.08, 115.25, 67.65, 42.79, 42.62, 21.29.

**HRMS (ESI+)** *m/z* calcd for $C_{26}H_{23}ClN_6O_3Na^+$ $[M+Na]^+$: 525.1418, found: 525.1458.

**Synthesis of IA-mTz-modified RS3$^{97\text{-}105}$ peptides**

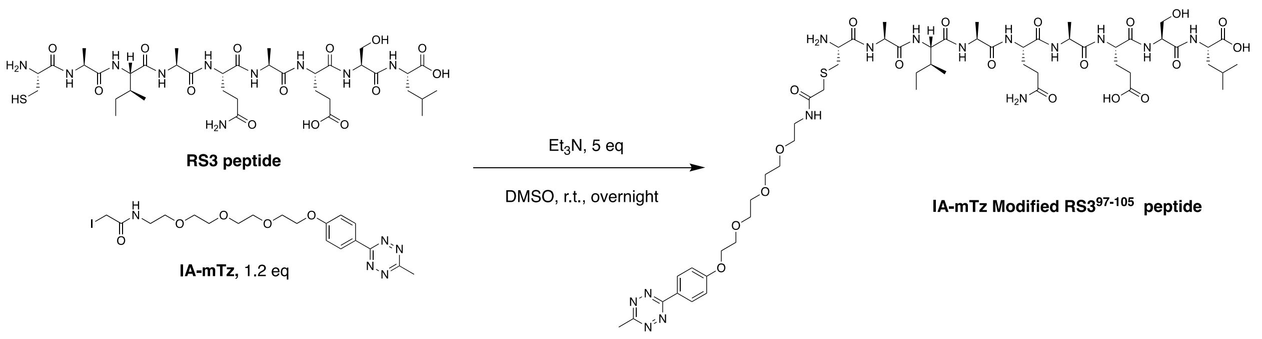

**RS3 peptide** (10 mg, 0.011 mmol, 1 eq, purchased from GenScript) and **IA-mTz** (7 mg, 0.0133 mmol, 1.2 eq) were dissolved in DMSO. Then $Et_3N$ (7.7 μL, 0.055 mmol, 5 eq)

were added and stirred in room temperature for overnight. The reaction was purified by prep-HPLC to provide IA-mTz-modified $RS3^{97-105}$ peptide as red solid. The HRMS mass peak of IA-mTz-modified $RS3^{97-105}$ peptide was shown at *m/z* 1308.6297 as $C_{56}H_{90}N_{15}O_{19}S^+$ $[M+H]^+$.

**[1]H NMR spectrum of KB02-mTz**

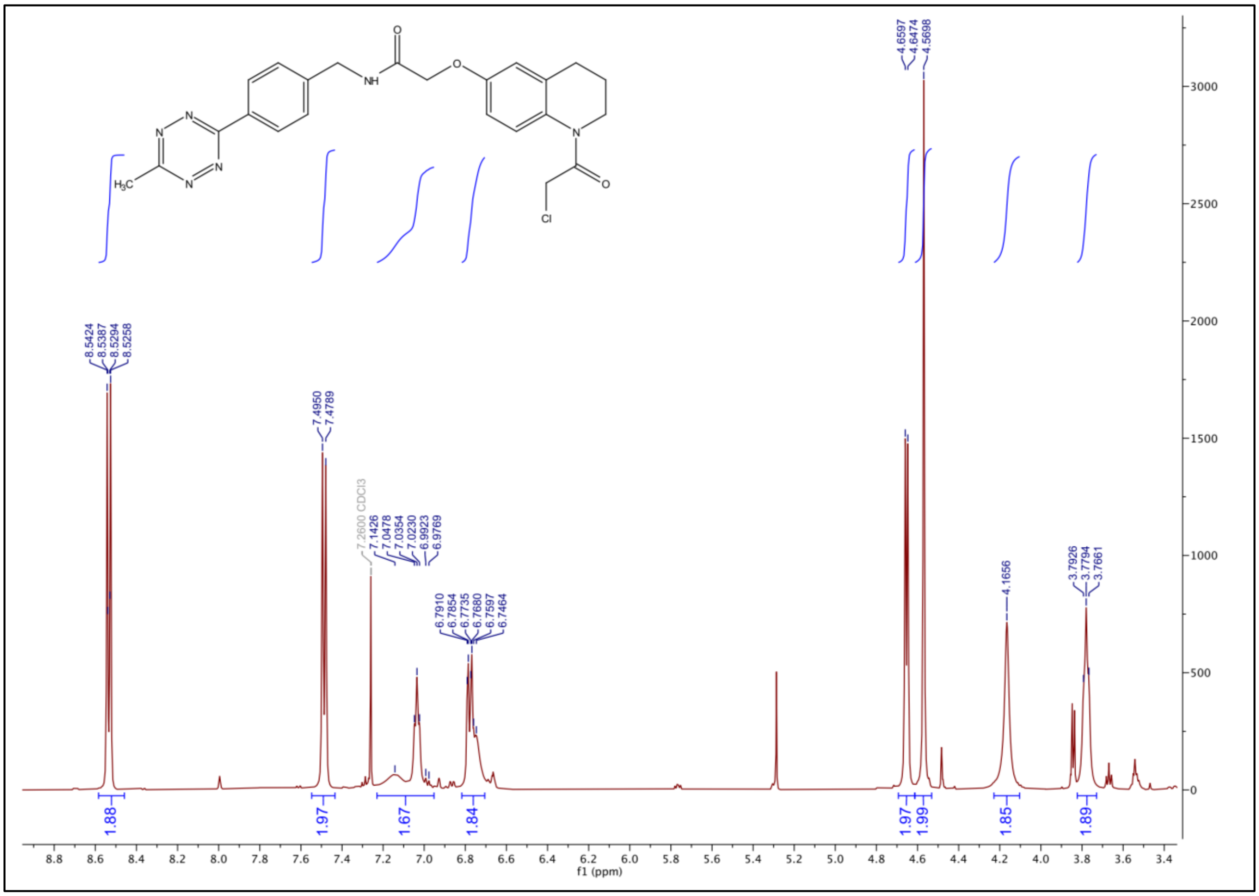

### $^{13}$C NMR spectrum of KB02-mTz

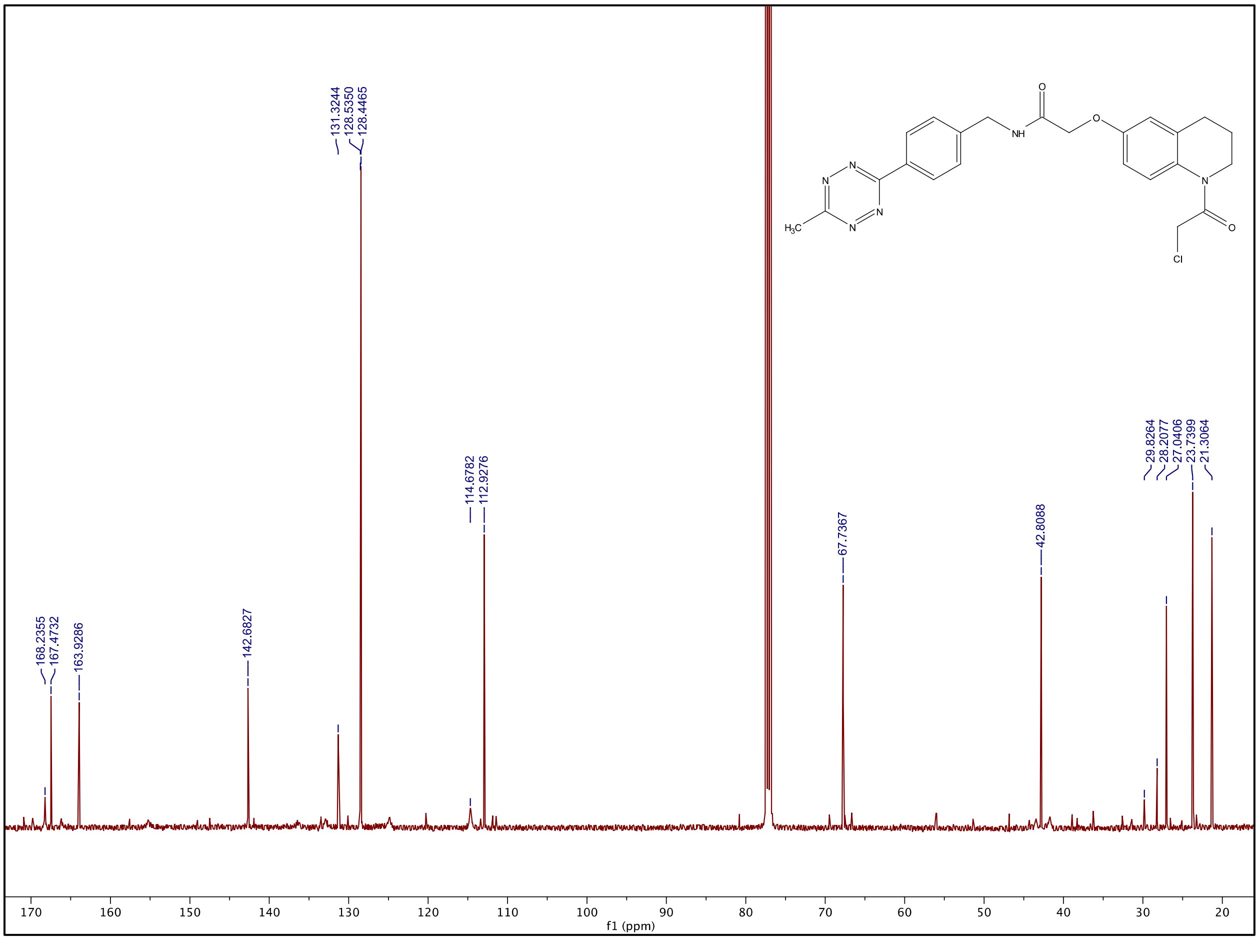

### [1]H NMR spectrum of KB05CAA-mTz

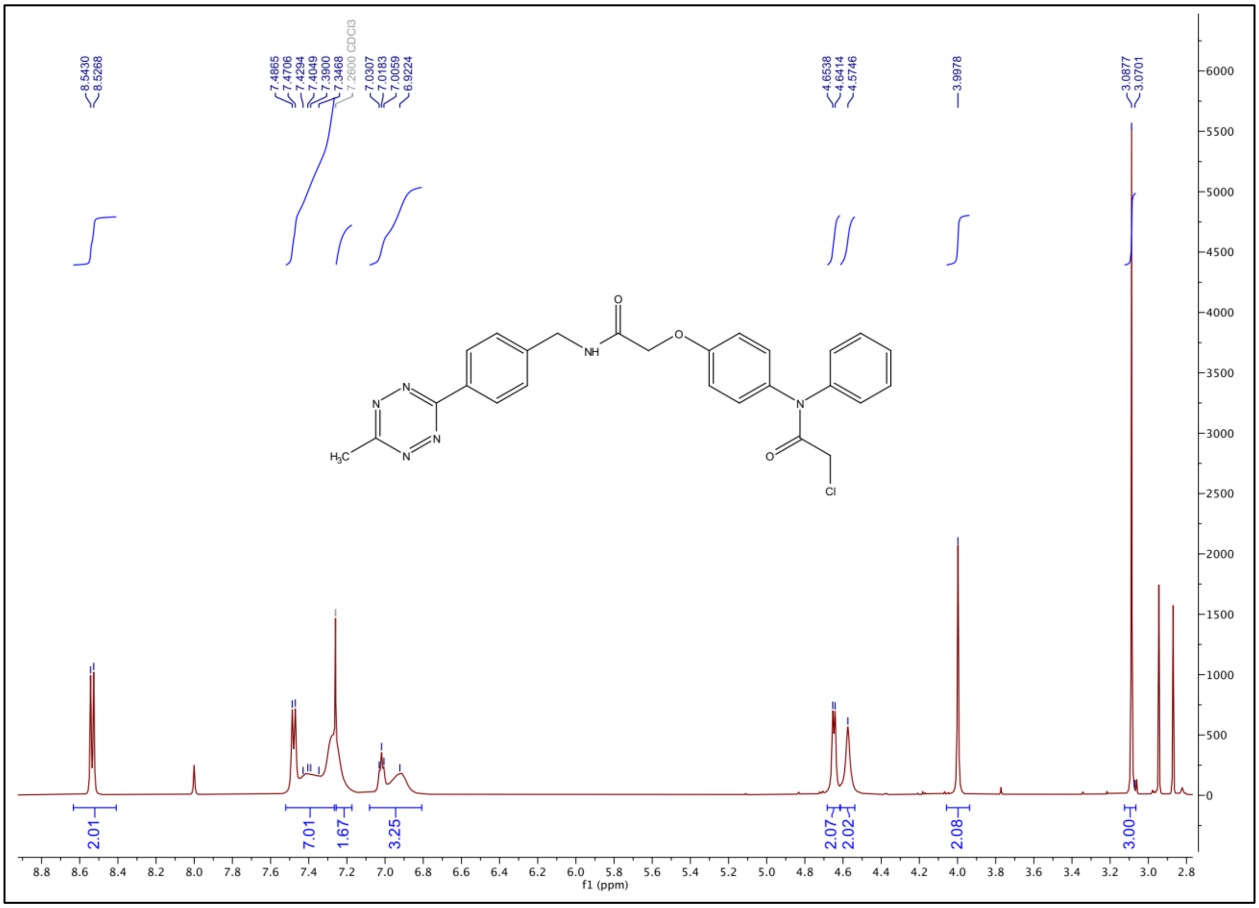

### $^{13}C$ NMR spectrum of KB05CAA-mTz

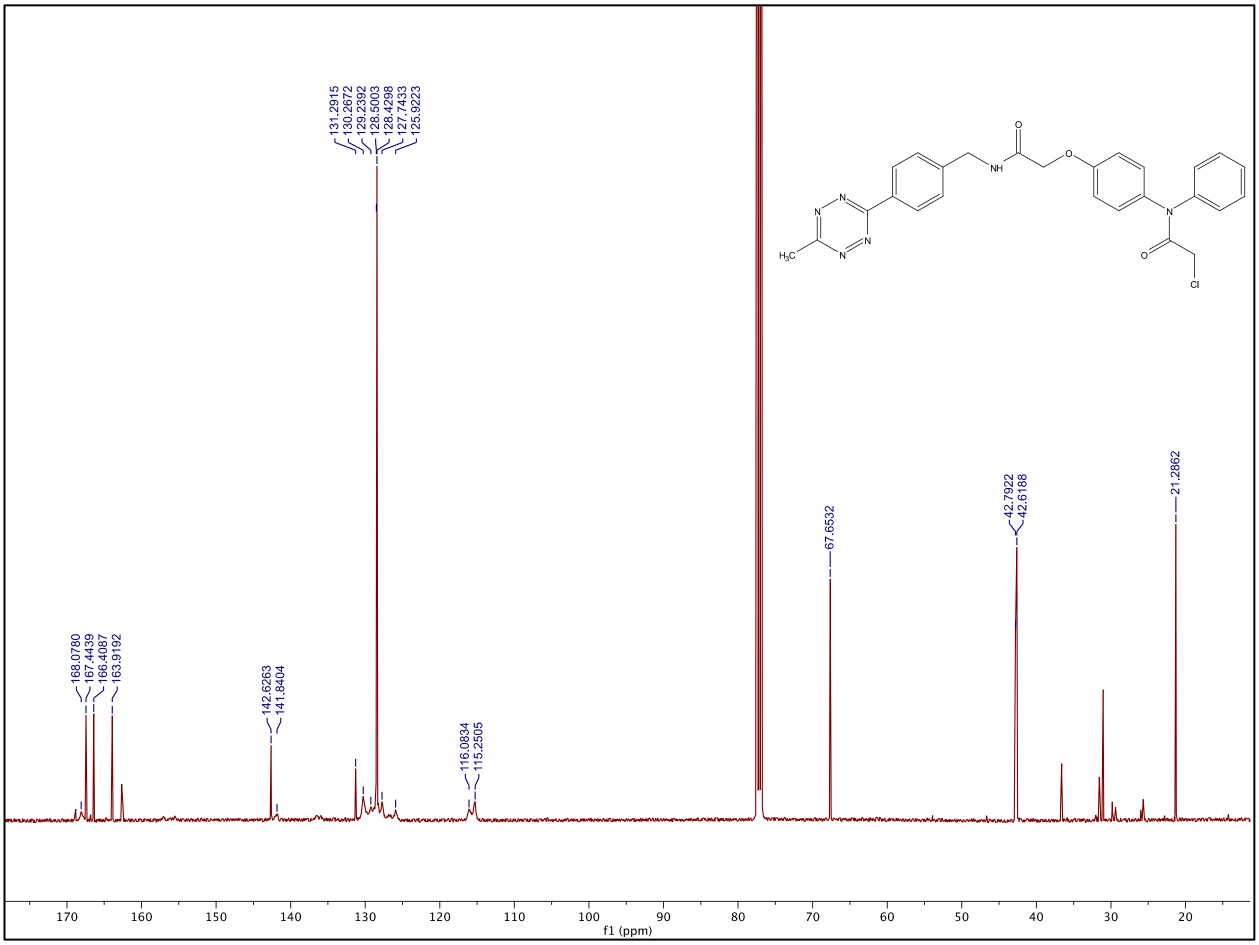
